# Dynamic changes of gut microbiota associated with blood glucose levels in sows from pregnancy through lactation

**DOI:** 10.1101/2022.03.08.483281

**Authors:** Mingjun Wang, Jiaju Li, Bo Chen, Haitao Li, Jun Ren

**Affiliations:** Center of Feed R&D Innovation, COFCO Feed Co., Ltd., Beijing 100005, China; Center of Biotechnology, COFCO Nutrition and Health Research Institute, Beijing 102209, China; Center of Animal Nutrition and Feed, COFCO Nutrition and Health Research Institute, Beijing 102209, China

**Keywords:** gut microbiota, blood glucose, sows, restricted feeding, pregnancy

## Abstract

Gut microbiota plays important roles in metabolism and physiological homeostasis throughout the lifespan of mammals, including in pregnancy. To evaluate the effects of practically adopted restricted feeding during gestation on blood glucose level and fecal microbiota composition, 31 sows were used to identify the shifts in bacterial assemblages by 16S rRNA sequencing in their early pregnancy, late pregnancy and late lactation stages. We find that the blood glucose was elevated significantly (*p* < 0.001) in sows offered abundant feed in the late lactation as compared with those of the same cohort in the pregnancy stages when on limited feed. Five bacterial genera were differentially abundant among three stages, and the abundances of *Streptococcus* (*p*_*adj*_ < 0.001) and *unclassified_f_Lachnospiraceae* (*p*_*adj*_ < 0.001) were increased, whereas the abundances of *Lactobacillus* (*p*_*adj*_ = 0.035) and *Escherichia-Shigella* (*p*_*adj*_< 0.001) were decreased in the latter two stages. The genus *Prevotellaceae_NK3B31_group* (*p*_*adj*_ = 0.026) displayed higher abundance in the late lactation stage. A significant correlation in the Pearson’s analysis indicates that three out of these five differentially abundant genera were related to blood glucose change. Our data show that restricted-fed gestating sows have aberrant blood glucose level and may affect the intestinal microbiota.

## Introduction

Restricted post-mating feeding strategy of pregnant sows is commonly applied under modern swine breeding conditions for the purpose of good health and optimal reproductive performance (Dourmad et al. 1996; Meunier-Salaün et al. 2001; Peltoniemi et al. 2000). This direction is based on the fact that over-feeding during gestation causes excessive body weight gain and fatness at the end of pregnancy, followed by reduced feed intake and inordinate weight loss in the lactating period, and thus promotes the occurrence of locomotion, farrowing and lactation problems (Dourmad et al. 1994; Lawlor et al. 2007; Thaker & Bilkei 2005). In Chinese swine feeding practices, hyperprolific sows are given access to a limited daily feed amount of 2-4 kg/sow from the start of the gestation till parturition, upon when they are fed *ad libitum*, with average daily feed intake reaching about 6 kg. On the contrary, it has been found that implementing restricted nutrient intake induces stereotypic behavior and fluctuates interprandial blood glucose and insulin levels in reproductive sows (de Leeuw et al. 2004), raising the consideration of insufficient feeling of satiation and potential adverse health effects.

Among the various life events, pregnancy is one of the most challenging times for the mammalian body (Koren et al. 2012). Over the course of gestation, the body undergoes substantial physiological changes including gut microbial diversity, contributing metabolic responses in return through intestinal host-microbial interactions (Nicholson et al. 2012). The mammalian gastrointestinal tract is home to myriads of bacteria and the gut microbiota plays a significant role in modulating intestinal morphology, food digestion, immune response, glucose homeostasis and health of the host (Ashida et al. 2011; Greiner & Bäckhed 2011; He et al. 2017). Changes in the gut microbiota are especially suggested to have connection with incidence of gestational diabetes mellitus, a symptom of hyperglycemia occurring during pregnancy (Angueira et al. 2015). The gut microbes are dynamic and evolve within a lifespan of the host (Yatsunenko et al. 2012). Accordingly, understanding the environmental factors that can shape the gut microbial community is crucial to gain insight into how this consortium of microbes is assembled and perturbed (Isaacson & Kim 2012; Korpela et al. 2016). It is well known that dietary changes can put impact on intestinal bacterial communities and their gene richness through modulating the supply of substrates for microbial growth and also the gut environment (Cotillard et al. 2013; Flint et al. 2017), but there are no existing studies describing the development of the gut microbiota in pregnant sows under restricted feeding.

In this study, multiparous sows were followed from pregnancy to one month postpartum to investigate the influence of long-term restricted feeding during pregnancy on blood glucose level and gut microbiota composition. Using 16S rRNA gene sequencing, we characterized bacterial taxa that were associated with alterations of the blood glucose profiles.

## Material and Methods

### Animals and sample collection

All procedures involving animals were carried out according to experimental protocols approved by the Animal Ethics Committee of COFCO Nutrition and Health Research Institute. 31 Landrace × Large White sows (artificially inseminated with semen from one Duroc boar) from the high health status herd at the Shunyu Pig Farm (Chengdu, China) were used in this study, with full approval from the farm owner. These animals were of similar age, parity and body conditions, and received no antibiotics or other medication. The herd had a history of vaccination against porcine reproductive and respiratory syndrome virus, parvovirus, pseudorabies virus, epidemic encephalitis B, and classical swine fever.

All of these sows were fed the same commercial sow pregnancy diet at 2.0 kg/d at the beginning of the study, and the feeding amount was progressively elevated to 3.0 kg/d before farrowing. After these sows gave birth to piglets and start lactating, free access was allowed to the same commercial lactation diet during the 4 weeks of lactation (17-20 weeks after mating). Details on sow diet ingredients and nutrient composition are provided in Table S1. Water was supplied under *ad libitum* strategy. The sows were returned to the general population of the farm after this study.

Fresh fecal samples were collected at the end of 4, 13, and 20 weeks after mating to represent early pregnancy, late pregnancy and late lactation stages of sows, respectively. An 8-hour fasting was executed before sampling. Approximately 200 g of each sample was individually harvested using Stool Collection Tubes (Stratec Biomedical, Birkenfeld, Germany) and then stored according to the manufacturer’s instructions before DNA extraction.

### Blood glucose value measurement

Preprandial blood samples taken from sows were obtained 2 h prior to feeding in the morning on the same days when collecting fecal samples. Blood glucose was determined by a handheld glucometer (Johnson & Johnson, New Brunswick, USA) using a small drop of blood through tail vein puncture modified from a previously reported protocol (Sanchez et al. 2016).

### Microbiome analyses

Bacterial genomic DNA of each fecal sample was extracted using the QIAamp Fast DNA Stool Mini Kit (Qiagen, Hilden, Germany). DNA concentration and purity was monitored by agarose electrophoresis. The V3-V4 region of the 16S rRNA bacterial gene (338-806) was then amplified using specific primers with the barcodes. Purified amplicons from each sample were pooled in equimolar amounts and paired-end sequenced (2 × 300 bp) on the Illumina MiSeq platform according to the standard protocols. The raw reads were deposited into the NCBI Sequence Read Archive (SRA) database (Accession Number: SRP117786).

Raw sequences were demultiplexed and quality-filtered using QIIME (version 1.17, http://qiime.org) (Caporaso et al. 2010) to eliminate all low quality sequence reads with the following criteria: 1) a minimum quality score of 20, 2) a minimum sequence length of 50 bp, 3) exact barcode matching and a maximum of two mismatches in the primer sequences, 4) no ambiguous bases in the sequences. Only reads that overlap longer than 10 bp were assembled otherwise discarded. The resulting trimmed sequences were then grouped into operational taxonomic units (OTUs) with 97% identity threshold using the USEARCH software based on the UPARSE algorithm (version 7.1, http://drive5.com/uparse) while chimeric sequences were identified and removed using UCHIME (Edgar et al. 2011). The taxonomy of each 16S rRNA gene sequence was analyzed by the RDP Classifier (http://rdp.cme.msu.edu) against the Silva (SSU128) 16S rRNA database using confidence threshold of 70%.

### Statistical analyses

Data analyses were performed with SPSS Statistics software (version 24.0, IBM Corporation, Armonk, USA) and R (version 3.3.1, http://www.r-project.org). Differences in blood glucose level of breeding sows among early pregnancy, late pregnancy and late lactation stages were assessed by use of one-way repeated measures ANOVA with Bonferroni post hoc test. Differences in α-diversity for the fecal microbiota were tested using Friedman test. Differences in microbiota composition between stages as assessed by β-diversity metrics were tested using Adonis (Anderson 2001) (999 permutations) at OTU level in the R vegan package (version 2.4-4, http://CRAN.R-project.org/package=vegan). Differences in abundance of taxa were tested using Kruskal-Wallis H test with *p*-values adjusted (*p*_*adj*_) for multiple testing by the FDR procedure, repeated measurements with Tukey post hoc test. Correlation between genera abundances and blood glucose was tested using Pearson’s *r*. Differences in value of Kyoto Encyclopedia of Genes and Genomes (KEGG) pathways among groups were tested by Kruskal-Wallis H test.

## Results

### Blood glucose level elevated significantly after restricted-fed gestating sows entered into the late lactation stage

31 multiparous sows were selected and monitored for blood glucose periodically after breeding with similar parities and body weights. The quantity of commercial feed supply was limited to 2 kg/sow per day at the beginning of pregnancy, and elevated gradually to 3 kg/sow per day before farrowing. Blood glucose was measured at 2-month intervals in the early pregnancy, late pregnancy and late lactation, respectively. Statistical analysis of the data shows that fasting concentrations of blood glucose over the gestation period did not differ between early and late pregnancy stages (*p* > 0.999), with concentrations of 3.27 ± 0.19 and 3.41 ± 0.13 mM, respectively (Fig. 1a). After litters were born and these sows started nursing, restricted feeding was abandoned and the glycemic average of sows before weaning rose to 4.57 ± 0.19 mM (*p* < 0.001). We also compared the changing pattern of the blood glucose of the sows under different feeding conditions, and the results demonstrate that using gestation crate or feeding station during the pregnancy caused a similar profile of blood glucose levels (Fig. 1b). These data collectively suggest that restricted feeding of gestating sows was accompanied by a relatively low blood glucose level in comparison to that of the late lactation stage.

**Figure 1.**
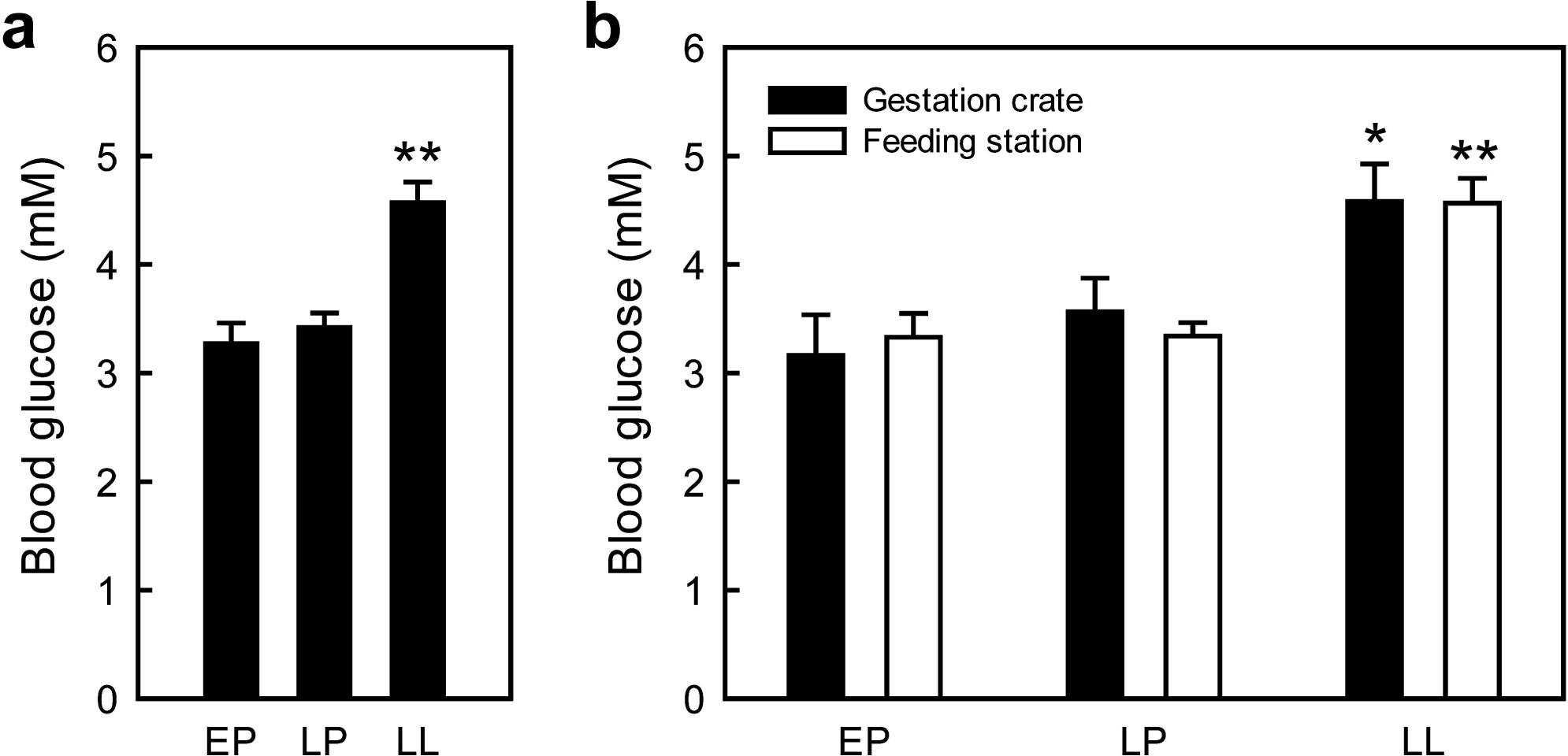
Blood glucose levels of breeding sows under restricted feeding in the early pregnancy, late pregnancy and late lactation stages. (a) Mean blood glucose level was significantly higher for sows in the late lactation than in pregnancy stages (*n* = 31). (b) Pregnant sows housed in a gestation crate (*n* = 11) or at a feeding station (*n* = 20) followed a similar pattern of blood glucose variation across time. Values are expressed as mean ± SEM. *, *p* < 0.05; **, *p* < 0.01 as compared with the other two stages (a) or the stage showing the lowest level (b). EP, early pregnancy; LP, late pregnancy; LL, late lactation.

### Sequencing data, OTUs and alpha diversity

We employed a culture-independent approach to compare the intestinal microbiota of the restricted-fed sows during pregnancy and lactation. Fresh fecal samples were collected on the same days of the blood sampling to perform the sequencing study. A total of 5,696,378 paired-end raw reads of 300 bp length were acquired by 16S rRNA sequencing after quality filtering, with 1,868,280, 1,906,196, and 1,921,902 reads for the early pregnancy, late pregnancy, and late lactation stages respectively. The total read length was 2.49 gigabases. Following subsampling, OTU picking and chimera checking, 3,201,897 clean sequences were obtained for downstream analysis and clustered into 2,420 non-singleton OTUs based on 97% sequence identity. Each sample had 1,031 OTUs on average (Table S2). 2,028 core OTUs were shared among the three stages studied, comprising approximately 84% of the total OTUs, while 27, 22, and 53 OTUs were uniquely identified at the early pregnancy, late pregnancy, and late lactation stages respectively (Fig. 2a).

**Figure 2.**
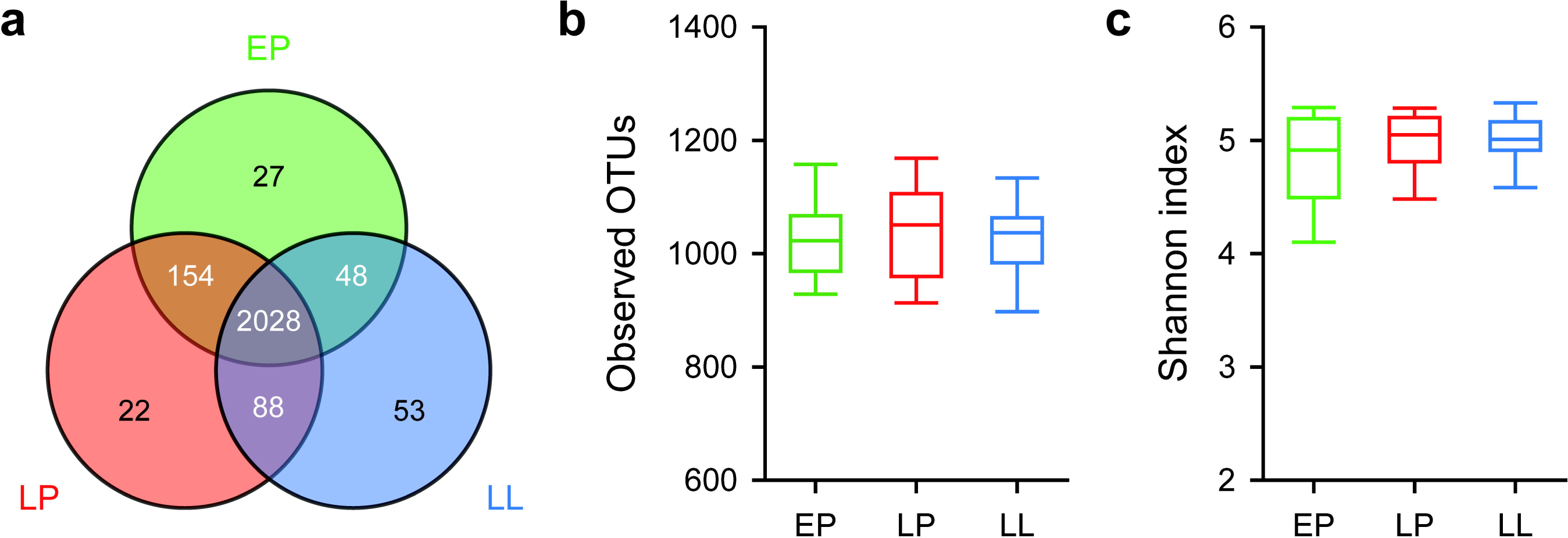
Comparison of the gut bacterial community composition, richness and diversity of restricted-fed sows at the early pregnancy, late pregnancy and late lactation stages. (a) Venn diagram was generated to illustrate overall overlap of OTUs among the three stages. (b) The bacterial abundance was represented by the numbers of OTUs. (c) Shannon index was used to estimate the diversity of the gut microbiota of the sows.

The values of Good’s coverage were 99.27%, 99.27% and 99.33% for the early pregnancy, late pregnancy, and late lactation stages respectively, indicating sufficient sequencing depth for further investigation of gut microbiota of restricted-fed gestating sows and capturing the dominant phylotypes. The OTU numbers and Shannon values of bacterial communities demonstrate that no significant differences were observed in the abundance (*p* = 0.597) and diversity (*p* = 0.206) of the gut microbiota at the three stages (Fig. 2b and c).

### Differences in bacterial community compositions among the three stages of restricted-fed sows

Adonis analysis using unweighted UniFrac metric, combined with principal coordinate analysis (PCoA), was employed to facilitate a statistical comparison of overall patterns of gut microbial composition among different stages of restricted-fed gestating sows (Fig. 3a). It is found that the distance between the three stages was significantly different (*r*^2^ = 0.028, *p* = 0.023), which indicates that the variation in our data set could be primarily explained by the gestation stages. Adonis based on weighted UniFrac (*r*^2^ = 0.047, *p* = 0.021) and Bray-Curtis (*r*^2^ = 0.034, *p* = 0.016) distances also showed a weak but significant contribution of physiological stage for shaping bacterial communities (Fig. S1). These results demonstrate that the association between stage and gut microbiota is driven by both the presence status and abundance of some taxa. Next, microbial composition analysis was performed to identify the main taxa contributing the difference between stages. The results illustrated in Fig. 3B describe the distribution of DNA sequences into phylum, and a total of 24 known bacterial phyla were shared by the three pregnancy/lactation-stage groups. *Firmicutes* and *Bacteroidetes* were the most dominant two phyla in all samples, regardless of stage, and comprised more than 85% of the total sequences. The bacterial abundances of distinct phyla differed in the three stages. *Firmicutes* (*p*_*adj*_ = 0.349) was the most prevalent phylum at early pregnancy, accounting for approximately 56% of the sequences. A relatively higher percentage (61% and 60%) of the sequences was assigned to *Firmicutes* at late pregnancy and late lactation stages. *Bacteroidetes* (*p*_*adj*_ = 0.583) was the second largest phylum in all groups, comprising approximately 24%, 26% and 27% in the early pregnancy, late pregnancy and late lactation groups, respectively. Among the other phyla, it is found that the bacterial abundances of three phyla revealed significant differences across the three stages (Fig. 3c-e). The proportions of bacteria belonging to *Proteobacteria* (*p*_*adj*_ < 0.001), *Verrucomicrobia* (*p*_*adj*_ = 0.017) and *Lentisphaerae* (*p*_*adj*_ < 0.001) decreased with the stages of pregnancy.

**Figure 3.**
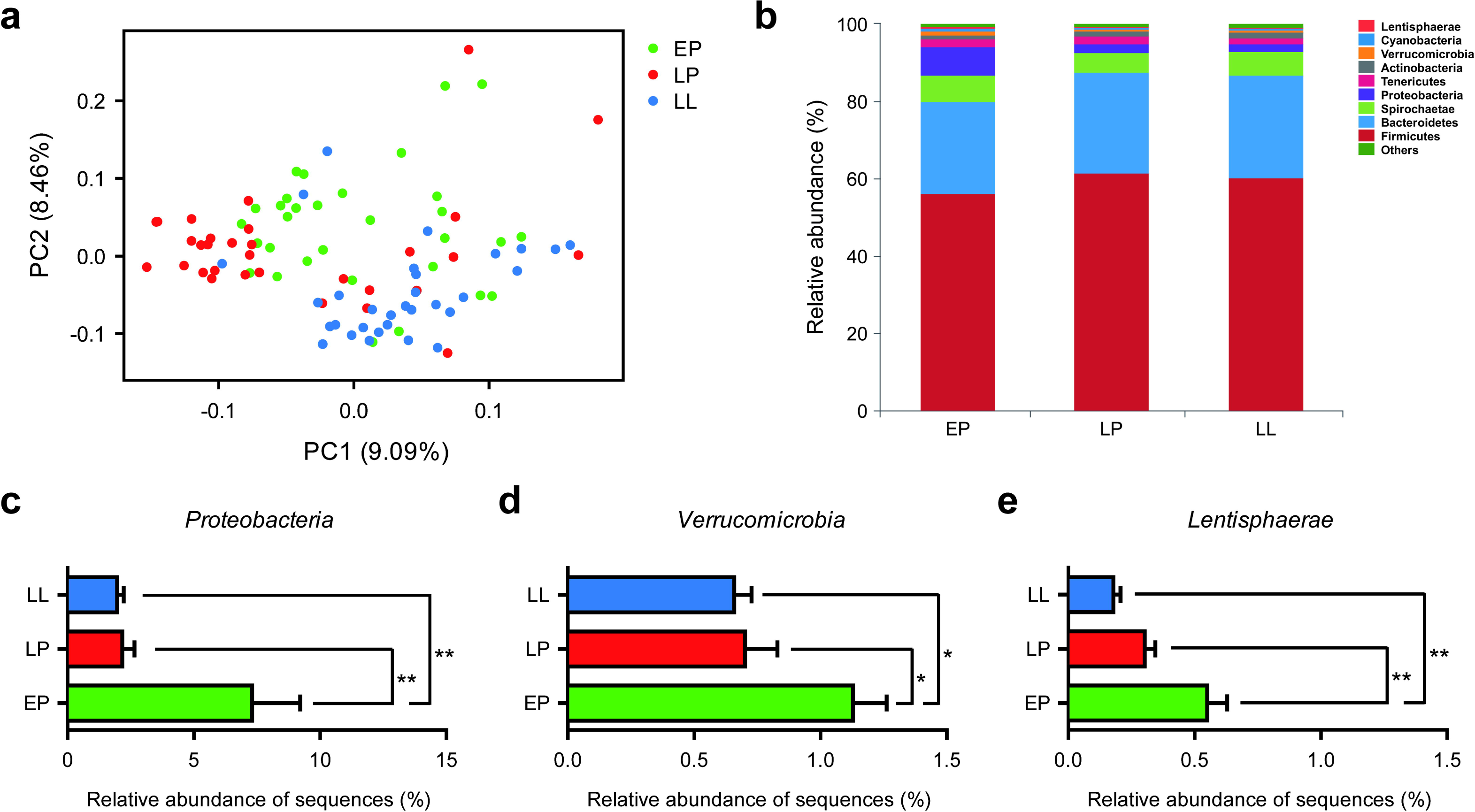
Taxonomic differences of gut microbiota among the early pregnancy, late pregnancy, and late lactation-stage groups. (a) PCoA plot based on unweighted UniFrac distance at OTU level reveals a gradient structure of taxonomic composition in which early pregnancy (green) occupies the intermediate position between late pregnancy (red) and late lactation (blue) samples. The percentage of variation explained by the principal coordinates is indicated on the axes. (b) Bacterial phylum distribution in fecal samples characterized by the average relative abundance of identified OTUs in individual groups. The top 10 abundant phyla are shown. (c-e) The relative abundance of three phyla, *Proteobacteria, Verrucomicrobia* and *Lentisphaerae*, significantly differed across groups. Values are expressed as mean ± SEM. *, *p* < 0.05; **, *p* < 0.01.

We also examined the compositional differences of bacterial communities by the other degrees of taxon. At the genus level, a total of 468 genera were identified from all samples, and 365 genera were shared by the three different groups. *Streptococcus* and *Lactobacillus* were the most predominant genera, accounting for 7.3% and 6.3% of total sequences, respectively (Fig. 4a). The bacterial abundance of five genera was found to be significantly different among three stages. The early pregnancy group had lower relative abundances of *Streptococcus* (Fig. 4b, *p*_*adj*_ < 0.001) and *unclassified_f_Lachnospiraceae* (Fig. 4c, *p*_*adj*_ < 0.001), and higher relative abundances of *Lactobacillus* (Fig. 4d, *p*_*adj*_ = 0.035) and *Escherichia-Shigella* (Fig. 4e, *p*_*adj*_ < 0.001) compared with the late pregnancy and late lactation groups, whilst the relative abundance of the genus *Prevotellaceae_NK3B31_group* (Fig. 4f, *p*_*adj*_ = 0.026) in late lactation was significantly higher than those in the other two earlier stages. In addition to the phylum and genus, bacterial abundance was also analyzed at the order level, and the relative abundance of *Bifidobacteriales* of restricted-fed sows in late lactation stage was found to be higher than those of the early and late pregnancy stages (Fig. S2, *p*_*adj*_ < 0.001).

**Figure 4.**
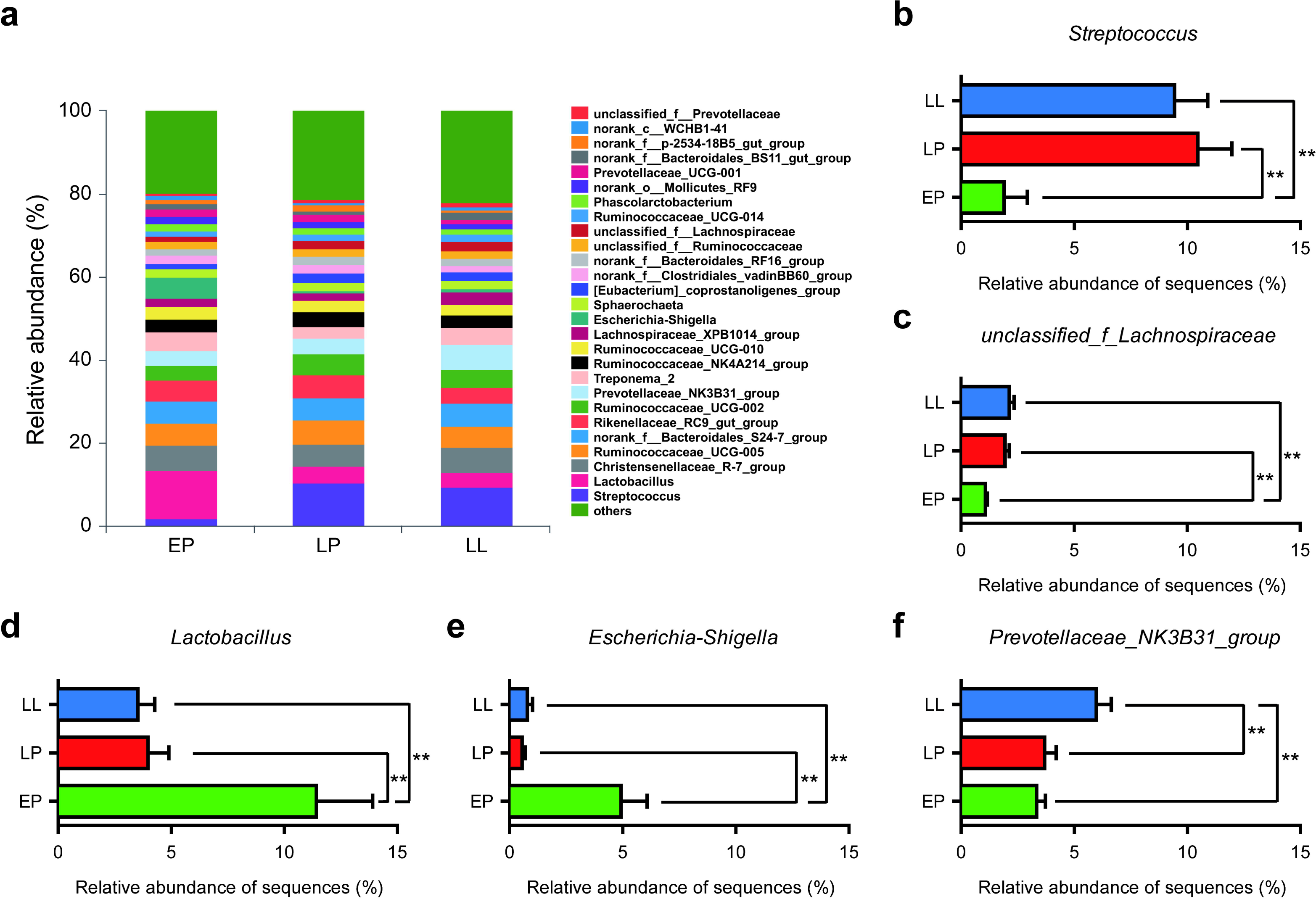
Bacterial composition significantly differed in accordance with the pregnancy/lactation stages of restricted-fed sows. (a) Distribution of the genera determined by an average percentage of the total number of bacterial 16S rRNA sequences in different stage groups. Only genera present in at least 1% of the samples are shown separately. (b-f) Comparison of the relative abundance of five genera, *Streptococcus, unclassified_f_Lachnospiraceae, Lactobacillus, Escherichia-Shigella* and *Prevotellaceae_NK3B31_group*, among the three stages. Values are expressed as mean ± SEM. **, *p* < 0.01.

### Correlation between the blood glucose level and the gut microbiota

We studied the correlation between genera whose relative abundances were higher than 1% and blood glucose of restricted-fed gestating sows. Notably, the abundances of *Prevotellaceae_NK3B31_group* (*r* = 0.209, *p* = 0.044) and *unclassified_f_Lachnospiraceae* (*r* = 0.208, *p* = 0.045) were positively correlated with fasting level of glucose in blood, and the abundance of the genus *Lactobacillus* (*r* = -0.227, *p* = 0.029) was negatively correlated with blood glucose level (Fig. 5a).

**Figure 5.**
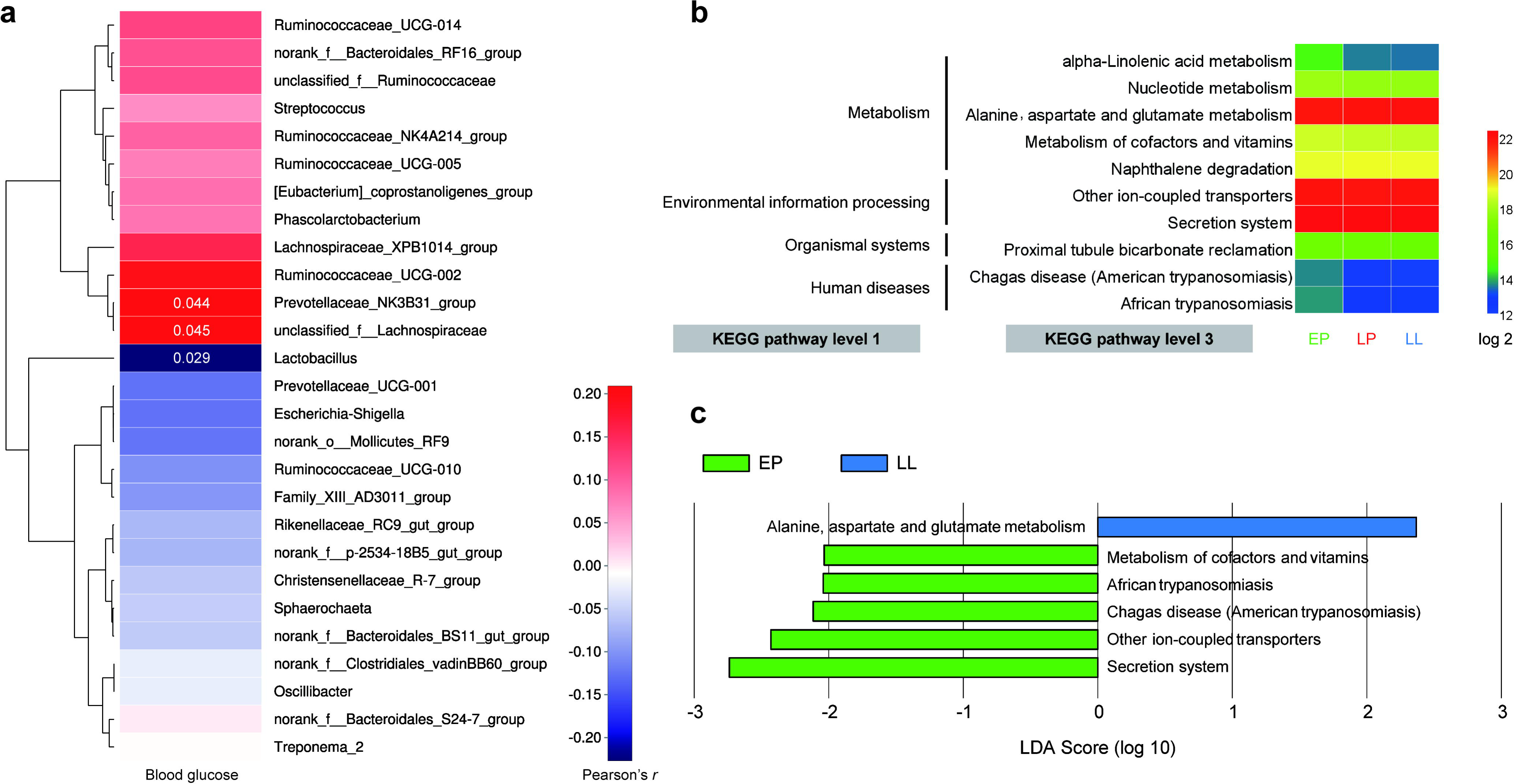
Correlation and metabolic implication of fasting blood glucose levels of restricted-fed gestating sows on the gut microbiota profiles. (a) Association between genera whose relative abundances were higher than 1% and blood glucose values. The colour key indicates Pearson’s *r* and the numbers in the cells represent *p* < 0.05. (b) Predicted function of gut microbiota among three stages of restricted-fed sows. The pathway abundances of samples within the same group were pooled and logarithmically transformed. Only functional features with *p* < 0.05 are shown using Kruskal-Wallis H test. (c) KEGG pathways significantly differentiated between the early pregnancy and late lactation stages identified by LEfSe with cutoff of LDA > 2.

We next used a computational tool, Phylogenetic Investigation of Communities by Reconstruction of Unobserved State (PICRUSt) (Langille et al. 2013), to explore the functional profiles of the sow gut microbiota and its association with metabolic regulation. The total 2,420 OTUs were normalized by its copy number and their metagenomic contributions were predicted from the KEGG pathways. In comparison between three stage groups, 10 pathways referred to as metabolism, environmental information processing, organismal system and human diseases were observed significantly different (Fig. 5b). Linear discriminant analysis coupled with effect size (LEfSe) (Segata et al. 2011) was further performed to identify pathways explaining the differences in each group. Five KEGG pathways were overrepresented in the early pregnancy stage, and one pathway, “alanine, aspartate and glutamate metabolism”, was overrepresented in the late lactation stage (Fig. 5c).

## Discussion

We have shown that gestating sows under restricted feeding experience recurrent hypoglycemia before parturition, and observed compositional and functional alterations of intestinal microbiota compared with when these sows enter into lactation. Our results suggest that albeit sharing most of the total 2,420 OTUs, several bacterial taxa including *Streptococcus, Lactobacillus, Prevotellaceae_NK3B31_group, Escherichia-Shigella*, and *unclassified_f_Lachnospiraceae* were differentially abundant among the early pregnancy, late pregnancy and late lactation stages.

Pregnancy is a time of dramatic host physiological changes and environmental remodeling. Previous reports about effects of gestation/lactation and feeding level on glucose metabolism in sows have been contradictory. Some studies showed that fasting glycemia did not change with stage of pregnancy and the higher feeding level (4 kg/d) decreased or did not affect plasma glucose concentration in restricted sows (Gatford et al. 2003; Père et al. 2000), and others indicated a lower basal plasma glucose level during lactation (Père & Etienne 2007). Very recently, a study in multiparous sows with free access to diets shows that the overall mean levels of preprandial blood glucose were similar between pregnancy and postpartum (Cheng et al. 2018). It is possible that these discrepancies are partly due to varying feeding strategy, and we assume that increased concentration of circulating glucose after farrowing, suggesting a state of insulin resistance, is more favored to milk production (Quesnel & Prunier 1998; Rehfeldt et al. 2004).

Several studies have reported that the diversity and composition of gut microbiota could be altered by nutritional and pregnancy status in other species (Jost et al. 2014; Queipo-Ortuño et al. 2013). Similarly, our PCoA analysis revealed a clustering of bacterial community structures according to stages of gestating sows under restricted nutrient intake, although we could poorly relate changes in alpha diversity indices to their pregnancy stages. It is interesting to note that nearly 84% of the total OTUs existed in the three stages that we examined, indicating a positive contribution of taxa abundance to the observed bacterial community shift.

Swine provide an attractive model and are utilized in experiments for human metabolic disease studies, given physiological and compositional similarity between these two organisms (Frese et al. 2015). Taken together with previous studies (Jost et al. 2014; Kim et al. 2015; Koren et al. 2012), our data support the assertion that *Firmicutes* represents the dominant phylum in the gut microbiota of pregnant mammals, including both swine and humans. In fact, *Firmicutes* was also found to be one of the most abundant groups at other growth stages of swine (Kim et al. 2011; Kim et al. 2015). Additionally, we observed a different changing pattern of the relative abundances of the phylum *Proteobacteria* with gestational age between swine and human. A prospective cohort study of Finnish mothers demonstrates that pregnant women showed an enrichment of *Proteobacteria* in late pregnancy (Koren et al. 2012), whereas we describe here a significant decrease in the relative abundances of this phylum on average in the perinatal period of sows. Considered in light of other findings indicating association of *Proteobacteria* with energy accumulation (Sun et al. 2016), our results reflect a high energy consumption state of hypoglycemic pregnant sows around farrowing. We also notice that one member of this group *Escherichia-Shigella* exhibited a parallel decline during the course of pregnancy and thus should be one of the potent candidates responsible for this community shift. Such facultative anaerobes contain large amounts of lipopolysaccharide and are active participants to induce metabolic inflammation, impaired glucose tolerance and development of type 2 diabetes (Kameyama & Itoh 2014; Khan et al. 2014).

*Streptococcus* and *Lactobacillus* are Gram-positive lactic acid bacteria belonging to the phylum *Firmicutes* and widely distributed in gastrointestinal tract. Similar to our finding, *Streptococcus* genus was also overrepresented in gravid women during the third trimester (33.72 ± 0.12 weeks) of pregnancy and 1 month postpartum, which is considered to improve commensal microbial sharing between mother and child and help educating the developing immune system (Koren et al. 2012). *Lactobacillus* has been linked to weight gain and obesity (Armougom et al. 2009), and is predominant in the vagina during pregnancy (Prince et al. 2015). Over the pregnancy course of restricted-fed sows the abundance of *Lactobacillus* was dramatically decreased and then remained relatively low during postpartum, whilst this tendency was significantly inversely correlated with blood glucose levels. In type 2 diabetes patients increased glycemic load has been proposed to promote the expansion of *Lactobacillus* and *Streptococcus* (Karlsson et al. 2013; Khan et al. 2014), reflecting a potential effect of the deficient blood glucose state on gut microbial alteration.

Two taxa that we found to differ among gestation/lactation stages were indicated to be positively associated with blood glucose concentration of restricted-fed sows. The genus *unclassified_f_Lachnospiraceae* refers to some unclassified bacterial species of the taxonomic family *Lachnospiraceae*. This group of microbes is abundant in the digestive tracts of mammals and many members are capable of producing short-chain fatty acids such as butyric acid (Meehan & Beiko 2014). Recently a specific *Lachnospiraceae* bacterium was identified to induce significant increases in fasting blood glucose levels and the development of obesity after colonization in an obese mouse model (Kameyama & Itoh 2014). Our observation about the elevated levels of *unclassified_f_Lachnospiraceae* during pregnancy suggest a compensatory response of gut microbes to the presence of hypoglycemia. The other differentially abundant taxon *Prevotellaceae_NK3B31_group* belonging to *Bacteroidetes* phylum has been reported to obviously decrease in type 2 diabetes rats (Wei et al. 2018). This is the only genus that we observed an enrichment in postpartum samples on average compared with those from stages of pregnancy. Such enrichment may reflect a rapid adaptation of the indigenous bacterial community to elevated blood glucose levels or otherwise a contribution to the metabolic and physiological requirements after parturition. The precise mechanisms underlying mutual interactions between these taxa and physiological circumstances including hypoglycemia and fetal development are still lacking and warrant further investigation.

Our PICRUSt analysis revealed a small but significant rise in numbers of predicted genes related to alanine, aspartate and glutamate metabolism in late lactation samples. This pathway includes synthesis of these amino acids from intermediates of glycolysis and citrate cycle, and allows the utilization of them for glucose generation under certain conditions. Increasing metabolism of these amino acids has been speculated to contribute to the development of insulin resistance (Sookoian & Pirola 2012), in accordance with our assumption regarding the elevated levels of fasting blood glucose in the late lactation stage.

In conclusion, the current data suggest pregnancy with restricted feeding to be associated with a marked lowering of the blood glucose level and a profound alteration of the gut microbiota of sows. Future metagenomic and metabolomics studies are needed to unravel the observed correlation between taxon abundance and blood glucose regulation in this study. Gaining detailed mechanistic insights of how the gut microbiota modulates glucose metabolism in sows may pave the way for novel interventions targeting metabolic regulation based on the microbiota.

## Acknowledgements

This work was supported by the Beijing Municipal Science & Technology Program (Z141100002614014), the National Key R&D Program of China (2018YFD0500600), and the Beijing Future Science & Technology Park Scientific Research Program. We thank Dr. Jian Xu (QIBEBT-CAS, Qingdao, China) for assistance with the trial design.

## Competing interests

The authors declare they have no competing interests.

## Additional files

Table S1. Ingredients and nutrient composition of sow pregnancy and lactation diets (as-fed basis).

Table S2. OTU table summary.

Figure S1. PCoA plot based on weighted UniFrac (a) and Bray-Curtis (b) distances at OTU level to represent taxonomic composition of early pregnancy, late pregnancy, and late lactation groups.

Figure S2. The relative abundance of the order *Bifidobacteriales* significantly differed between groups. **, *p* < 0.01.

## Notes

### Competing Interest Statement

The authors have declared no competing interest.

