## Supplementary material for "Dynamic changes of gut microbiota associated with blood glucose levels in sows from pregnancy through lactation": Table S1

**Table S1. Ingredients and nutrient composition of sow pregnancy and lactation diets (as-fed basis)**

|  | **Pregnancy diet** | **Lactation diet** |
| --- | --- | --- |
| **Ingredient, %** |  |  |
| Corn | 51.99 | 57.21 |
| Soybean meal, 46% CP | 12.70 | 20.80 |
| Wheat bran | 13.00 | 5.80 |
| Sugar beet pulp | 18.00 | 5.00 |
| Fat powder | 0.00 | 3.00 |
| Enzymolytic soybean meal | 0.00 | 2.00 |
| Soybean oil | 0.50 | 1.50 |
| Dicalcium phosphate | 0.85 | 1.35 |
| Calcium carbonate | 0.90 | 1.00 |
| Premix | 1.00 | 1.00 |
| Salt | 0.43 | 0.32 |
| L-lysine hydrochloride | 0.10 | 0.25 |
| Sodium bicarbonate | 0.10 | 0.20 |
| Choline chloride | 0.10 | 0.15 |
| Potassium chloride | 0.10 | 0.15 |
| Mildewcide | 0.10 | 0.10 |
| Composite antioxidant | 0.07 | 0.07 |
| L-threonine | 0.06 | 0.06 |
| Oregano oil | 0.00 | 0.02 |
| L-carnitine | 0.00 | 0.02 |
| **Nutrient composition** |  |  |
| Net energy, kcal/kg | 2247.82 | 2448.97 |
| Crude protein, % | 13.80 | 16.50 |
| Crude fat, % | 3.40 | 5.90 |
| Crude fiber, % | 6.22 | 3.78 |
| NDF, % | 19.86 | 13.02 |
| ADF, % | 7.68 | 4.79 |
| Lys, % | 0.75 | 1.05 |
| Ca, % | 0.89 | 0.89 |
| Total P, % | 0.50 | 0.57 |
