## Supplementary material for "Dynamic changes of gut microbiota associated with blood glucose levels in sows from pregnancy through lactation": Table S2

**Table S2. OTU table summary**

| **Sow ID** | **Group** | **Sample ID** | **Reads** | **OTUs** | **Good’s** | **ACE** | **Chao1** | **Shannon** | **Simpson** |
| --- | --- | --- | --- | --- | --- | --- | --- | --- | --- |
| Y29804 | EP | A1 | 56151 | 969 | 99.49% | 1097 | 1094 | 5.2546 | 0.0139 |
| R18 | EP | A2 | 60325 | 1106 | 99.15% | 1369 | 1401 | 4.3209 | 0.1074 |
| 727 | EP | A3 | 60855 | 1023 | 99.19% | 1277 | 1283 | 4.4951 | 0.0512 |
| E2 | EP | A4 | 50285 | 1014 | 99.36% | 1188 | 1207 | 5.1753 | 0.0149 |
| 1724 | EP | A5 | 52720 | 1001 | 99.23% | 1248 | 1256 | 4.8204 | 0.0379 |
| R20 | EP | A6 | 51384 | 1024 | 99.41% | 1171 | 1185 | 5.1763 | 0.0156 |
| E13 | EP | A7 | 50696 | 982 | 99.31% | 1194 | 1223 | 5.1579 | 0.0161 |
| E11 | EP | A8 | 70032 | 958 | 99.34% | 1161 | 1166 | 4.9018 | 0.0254 |
| J113 | EP | A9 | 65164 | 1170 | 99.05% | 1479 | 1543 | 4.5999 | 0.0571 |
| E14 | EP | A10 | 50923 | 1167 | 99.25% | 1372 | 1409 | 5.3442 | 0.0138 |
| E16 | EP | A11 | 57838 | 1121 | 99.10% | 1407 | 1445 | 4.2082 | 0.1077 |
| 230 | EP | B1 | 68098 | 925 | 99.33% | 1139 | 1150 | 4.8944 | 0.0245 |
| A4 | EP | B2 | 50530 | 968 | 99.29% | 1188 | 1219 | 4.9156 | 0.0220 |
| 80 | EP | B3 | 69560 | 944 | 99.40% | 1118 | 1113 | 5.1912 | 0.0124 |
| 391 | EP | B4 | 70450 | 1037 | 99.27% | 1255 | 1242 | 4.7284 | 0.0369 |
| 899 | EP | B5 | 53025 | 1023 | 99.27% | 1255 | 1272 | 4.4041 | 0.0769 |
| JY12 | EP | B6 | 69005 | 1061 | 99.15% | 1351 | 1378 | 4.9947 | 0.0243 |
| 726 | EP | B7 | 61064 | 1009 | 99.29% | 1210 | 1211 | 4.0769 | 0.1264 |
| 272 | EP | B8 | 58373 | 1058 | 99.31% | 1250 | 1277 | 5.2059 | 0.0144 |
| J181 | EP | B9 | 52682 | 1024 | 99.23% | 1273 | 1302 | 4.9198 | 0.0236 |
| 602 | EP | B10 | 69720 | 882 | 99.28% | 1114 | 1125 | 3.5395 | 0.1737 |
| X11 | EP | B11 | 54381 | 1081 | 99.29% | 1279 | 1311 | 5.2915 | 0.0129 |
| 984 | EP | B12 | 59946 | 905 | 99.20% | 1199 | 1176 | 2.9875 | 0.3182 |
| 454 | EP | B13 | 64665 | 942 | 99.32% | 1153 | 1147 | 4.7715 | 0.0355 |
| 4003 | EP | B14 | 69788 | 997 | 99.42% | 1160 | 1158 | 5.3896 | 0.0109 |
| J119 | EP | B15 | 65950 | 995 | 99.32% | 1184 | 1202 | 4.7015 | 0.0319 |
| 8128 | EP | B16 | 69430 | 1062 | 99.26% | 1284 | 1307 | 4.9198 | 0.0291 |
| 224 | EP | B17 | 71287 | 990 | 99.22% | 1249 | 1253 | 4.4310 | 0.0593 |
| J212 | EP | B18 | 55995 | 1178 | 99.21% | 1400 | 1424 | 5.2839 | 0.0185 |
| 997 | EP | B19 | 50770 | 1110 | 99.22% | 1342 | 1387 | 5.1633 | 0.0187 |
| J160 | EP | B20 | 57188 | 1067 | 99.26% | 1298 | 1326 | 5.2126 | 0.0178 |
| Y29804 | LP | C1 | 74381 | 905 | 99.40% | 1077 | 1103 | 4.8868 | 0.0272 |
| R18 | LP | C2 | 63445 | 1052 | 99.21% | 1303 | 1288 | 4.4073 | 0.0915 |
| 727 | LP | C3 | 62056 | 960 | 99.30% | 1166 | 1214 | 4.6997 | 0.0289 |
| E2 | LP | C4 | 64897 | 1085 | 99.31% | 1279 | 1300 | 5.3759 | 0.0116 |
| 1724 | LP | C5 | 62622 | 1055 | 99.26% | 1271 | 1295 | 5.0850 | 0.0196 |
| R20 | LP | C6 | 53228 | 947 | 99.39% | 1111 | 1135 | 4.7708 | 0.0268 |
| E13 | LP | C7 | 53127 | 907 | 99.34% | 1128 | 1128 | 5.1249 | 0.0133 |
| E11 | LP | C8 | 62177 | 991 | 99.36% | 1178 | 1189 | 5.2755 | 0.0117 |
| J113 | LP | C9 | 55172 | 1136 | 99.11% | 1437 | 1439 | 5.1915 | 0.0206 |
| E14 | LP | C10 | 63976 | 950 | 99.31% | 1166 | 1218 | 5.1303 | 0.0125 |
| E16 | LP | C11 | 55733 | 992 | 99.29% | 1204 | 1225 | 4.1040 | 0.1216 |
| 230 | LP | D1 | 73948 | 898 | 99.31% | 1115 | 1137 | 4.5262 | 0.0541 |
| A4 | LP | D2 | 69415 | 1173 | 99.17% | 1420 | 1468 | 5.0363 | 0.0329 |
| 80 | LP | D3 | 54714 | 1040 | 99.16% | 1344 | 1339 | 5.0403 | 0.0179 |
| 391 | LP | D4 | 62120 | 1170 | 99.16% | 1420 | 1472 | 5.0492 | 0.0277 |
| 899 | LP | D5 | 62320 | 1057 | 99.33% | 1246 | 1288 | 5.2012 | 0.0188 |
| JY12 | LP | D6 | 52731 | 1162 | 99.18% | 1413 | 1402 | 5.0025 | 0.0340 |
| 726 | LP | D7 | 57761 | 1056 | 99.33% | 1244 | 1216 | 4.9194 | 0.0332 |
| 272 | LP | D8 | 59314 | 1221 | 99.21% | 1442 | 1472 | 5.3414 | 0.0197 |
| J181 | LP | D9 | 61610 | 953 | 99.30% | 1164 | 1152 | 5.0228 | 0.0169 |
| 602 | LP | D10 | 54813 | 1008 | 99.33% | 1206 | 1215 | 5.0930 | 0.0214 |
| X11 | LP | D11 | 54622 | 998 | 99.34% | 1187 | 1203 | 5.2099 | 0.0133 |
| 984 | LP | D12 | 54785 | 1118 | 99.27% | 1323 | 1337 | 5.0979 | 0.0220 |
| 454 | LP | D13 | 66933 | 1000 | 99.30% | 1209 | 1216 | 4.8120 | 0.0321 |
| 4003 | LP | D14 | 74359 | 970 | 99.34% | 1165 | 1153 | 4.9252 | 0.0281 |
| J119 | LP | D15 | 53707 | 1094 | 99.30% | 1281 | 1304 | 5.1117 | 0.0248 |
| 8128 | LP | D16 | 57064 | 1090 | 99.18% | 1357 | 1354 | 5.2705 | 0.0129 |
| 224 | LP | D17 | 64224 | 1051 | 99.25% | 1284 | 1250 | 4.4706 | 0.0622 |
| J212 | LP | D18 | 71861 | 937 | 99.22% | 1216 | 1248 | 4.6563 | 0.0465 |
| 997 | LP | D19 | 67663 | 1127 | 99.22% | 1361 | 1361 | 5.2441 | 0.0158 |
| J160 | LP | D20 | 61418 | 1106 | 99.26% | 1319 | 1348 | 5.2872 | 0.0127 |
| Y29804 | LL | E1 | 60643 | 1060 | 99.37% | 1241 | 1234 | 5.3966 | 0.0118 |
| R18 | LL | E2 | 55377 | 1066 | 99.37% | 1236 | 1243 | 5.0790 | 0.0314 |
| 727 | LL | E3 | 65151 | 1021 | 99.33% | 1207 | 1227 | 5.0795 | 0.0181 |
| E2 | LL | E4 | 54221 | 1004 | 99.30% | 1217 | 1230 | 4.9130 | 0.0257 |
| 1724 | LL | E5 | 64567 | 924 | 99.28% | 1172 | 1175 | 4.9842 | 0.0163 |
| R20 | LL | E6 | 53173 | 960 | 99.40% | 1127 | 1150 | 4.9826 | 0.0305 |
| E13 | LL | E7 | 63080 | 1042 | 99.24% | 1274 | 1270 | 5.0040 | 0.0205 |
| E11 | LL | E8 | 66618 | 1028 | 99.36% | 1197 | 1214 | 4.5588 | 0.0572 |
| J113 | LL | E9 | 73166 | 1057 | 99.32% | 1250 | 1262 | 5.1897 | 0.0187 |
| E14 | LL | E10 | 60706 | 1018 | 99.37% | 1186 | 1215 | 5.3676 | 0.0102 |
| E16 | LL | E11 | 71189 | 1132 | 99.34% | 1305 | 1337 | 5.3201 | 0.0174 |
| 230 | LL | F1 | 55279 | 844 | 99.46% | 1002 | 987 | 4.9771 | 0.0164 |
| A4 | LL | F2 | 58010 | 1063 | 99.25% | 1281 | 1266 | 4.4231 | 0.0628 |
| 80 | LL | F3 | 57613 | 1050 | 99.25% | 1267 | 1253 | 4.7653 | 0.0241 |
| 391 | LL | F4 | 68116 | 1121 | 99.29% | 1308 | 1319 | 5.2577 | 0.0172 |
| 899 | LL | F5 | 70077 | 1169 | 99.24% | 1372 | 1394 | 5.0450 | 0.0298 |
| JY12 | LL | F6 | 61226 | 1123 | 99.28% | 1329 | 1342 | 4.9837 | 0.0481 |
| 726 | LL | F7 | 56398 | 1053 | 99.44% | 1188 | 1202 | 5.3319 | 0.0125 |
| 272 | LL | F8 | 54082 | 1173 | 99.27% | 1369 | 1401 | 5.1643 | 0.0330 |
| J181 | LL | F9 | 67197 | 961 | 99.36% | 1154 | 1157 | 5.0303 | 0.0163 |
| 602 | LL | F10 | 62102 | 997 | 99.37% | 1169 | 1161 | 4.9423 | 0.0212 |
| X11 | LL | F11 | 68813 | 1134 | 99.17% | 1386 | 1383 | 4.8556 | 0.0472 |
| 984 | LL | F12 | 51524 | 1015 | 99.29% | 1228 | 1206 | 4.6809 | 0.0318 |
| 454 | LL | F13 | 71344 | 984 | 99.37% | 1153 | 1178 | 5.1519 | 0.0142 |
| 4003 | LL | F14 | 57815 | 1037 | 99.20% | 1292 | 1351 | 5.0102 | 0.0223 |
| J119 | LL | F15 | 69032 | 1019 | 99.36% | 1187 | 1200 | 4.4346 | 0.0922 |
| 8128 | LL | F16 | 68918 | 880 | 99.47% | 1031 | 1021 | 4.9431 | 0.0237 |
| 224 | LL | F17 | 65494 | 891 | 99.40% | 1073 | 1133 | 5.3029 | 0.0094 |
| J212 | LL | F18 | 55149 | 1045 | 99.28% | 1255 | 1286 | 4.7137 | 0.0699 |
| 997 | LL | F19 | 65048 | 978 | 99.36% | 1153 | 1181 | 5.0760 | 0.0160 |
| J160 | LL | F20 | 50774 | 1045 | 99.32% | 1233 | 1284 | 5.1500 | 0.0176 |
