## Supplementary figures and images for "Dynamic changes of gut microbiota associated with blood glucose levels in sows from pregnancy through lactation"

### Figure S1

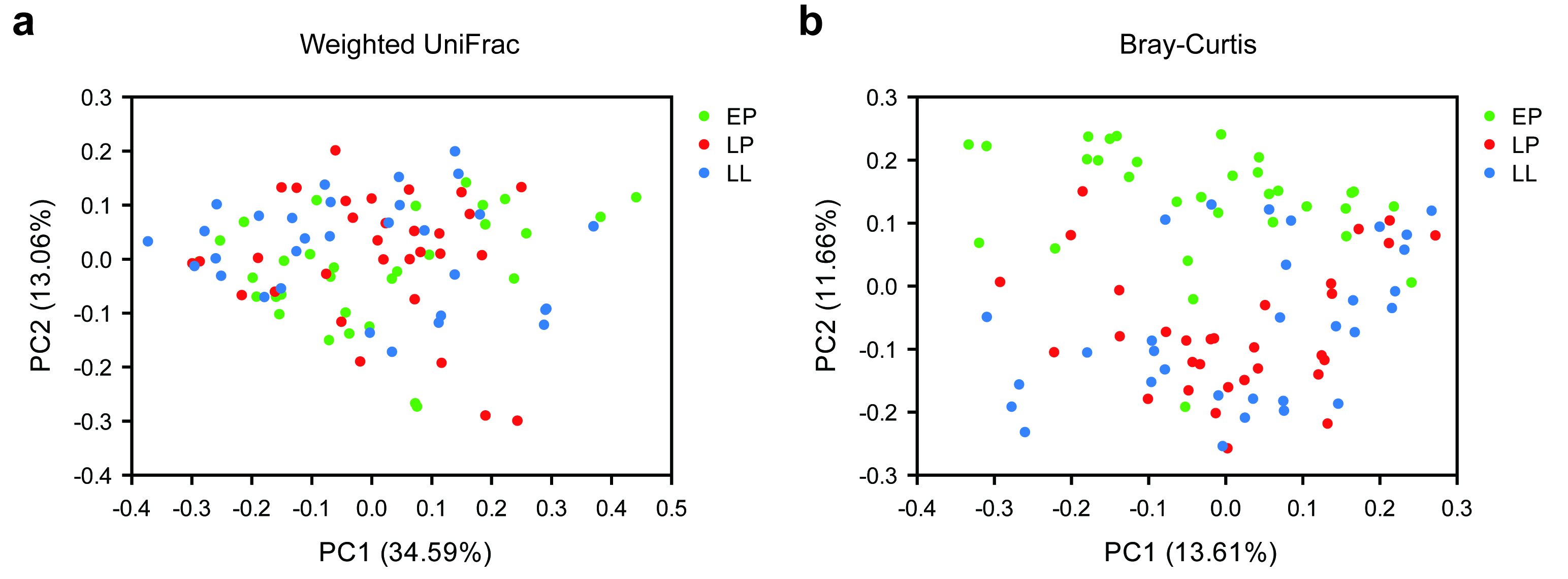

### Figure S2

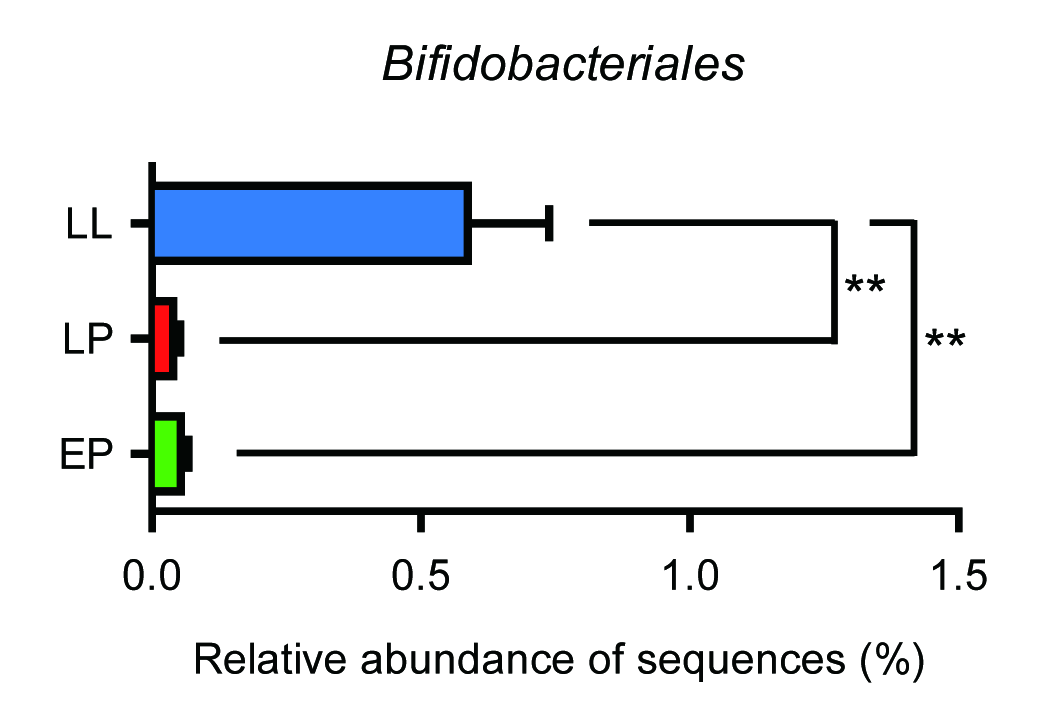
